# Mapping connectivity and conservation opportunity on agricultural lands across the conterminous United States

**DOI:** 10.1101/2022.10.08.511378

**Authors:** Justin P. Suraci, Caitlin E. Littlefield, Charlie C. Nicholson, Mitchell C. Hunter, Ann Sorensen, Brett G. Dickson

## Abstract

1. Depending on management practices, agricultural lands can either pose substantial barriers to the movement of native species or can support landscape connectivity by linking areas of high-quality habitat. Balancing connectivity and sustainable food production on agricultural lands is critical to conservation in the conterminous United States (CONUS) where agriculture makes up close to half of total land area. However, limited guidance exists on where to target conservation resources to maximize benefits for native species and food security.
2. To quantify the potential contribution of agricultural lands to the movement of organisms, we developed a novel method for estimating agricultural management intensity (based on remotely sensed temporal variation in vegetation cover on croplands and pastures) and incorporated these estimates into a CONUS-wide, circuit-theory based model of ecological flow connectivity. We then combined our connectivity results with data on the productivity, versatility, and resilience of agricultural lands (PVR) to identify conservation opportunities that support both biodiversity and food production.
3. The highest levels of connectivity on agricultural lands occurred on relatively unmodified rangelands and on cropland and pasture in close proximity to large amounts of natural land cover.
4. Mapping connectivity and PVR across CONUS revealed 10.2 Mha of agricultural lands (2.7%) with high value for both connectivity and food production, as well as large amounts of agricultural land (>140 Mha in total) with high value for either cultivation or supporting biodiversity (e.g., through ecological restoration).
5. Drawing on these findings, we provide recommendations on the types of conservation approaches most suitable for a given agricultural system and link these recommendations to specific government incentive programs. To help facilitate conservation planning based on our results, we have developed an interactive web application, allowing users to visualize the spatial data developed here within their regions of interest.

## Introduction

Preserving and enhancing the natural movement of organisms is critical to mitigating the current biodiversity crisis (Tilman *et al*. 2017) and is a key strategy for promoting species adaptations to climate change (Heller and Zavaleta 2009), with well-connected landscapes facilitating gene flow, migration, dispersal, and range shifts (McRae and Beier 2007; Littlefield *et al*. 2019). In the United States, private agricultural lands may play an important role in facilitating such ecological flows by providing linkages between areas of high-quality habitat (Kremen and Merenlender 2018; Garibaldi *et al*. 2021). Indeed, agricultural lands (including cropland, pasture, and rangeland) compose almost half the land area in the conterminous United States (CONUS) and, in many areas of the country, have continued to expand over the last decade (Lark *et al*. 2020). This trend is anticipated to continue (Sohl *et al*. 2014), underscoring the importance of centering agricultural landscapes in any comprehensive assessment of connectivity across the U.S.

Agricultural expansion, particularly high intensity crop production, has been a major driver of biodiversity declines globally through habitat loss, pesticide use, and the impacts of mowing and harvest (Newbold *et al*. 2015; Stanton *et al*. 2018). Intensively farmed areas may additionally represent substantial barriers to movement for a variety of taxa (Wimberly *et al*. 2018; Maas *et al*. 2021). However, low-intensity agriculture and wildlife-friendly management practices (e.g., grassland or forest strips, diversification of crops planted) can reduce these barriers to movement and even facilitate the flow of organisms across agricultural landscapes (Kremen and Merenlender 2018; Maas *et al*. 2021). Each year, governments spend billions of dollars globally to incentivize wildlife-friendly farming and other agri-environment schemes (Donald and Evans 2006), though limited information exists on where to target such financial incentives to maximize biodiversity benefits, potentially leading to the haphazard allocation of resources (Polasky *et al*. 2008; Kremen and Merenlender 2018).

Increases in global food production (of at least 25% by 2050; Hunter *et al*. 2017) will be necessary to support a growing human population. At the same time, climate change and urban and suburban expansion pose potential threats to food security by reducing the amount of land area that is highly suitable for cultivation (Tu *et al*. 2021; Kummu *et al*. 2021). It is therefore imperative to balance the dual goals of promoting biodiversity and safeguarding the working lands that are most critical for food production (Leclère *et al*. 2020). Two general strategies have been proposed for balancing biodiversity and agricultural objectives: ‘land sharing’, i.e., maintaining or enhancing the capacity of cultivated lands to support biodiversity through wildlife-friendly farming practices, potentially at the expense of yield; and ‘land sparing’, which advocates intensifying food production in some areas while preventing the expansion of agriculture into more natural landscapes, e.g., through formal protection (Fischer *et al*. 2008; Phalan *et al*. 2011; Grass *et al*. 2019). The feasibility and desirability of land sharing vs. sparing will depend on local context (e.g., biophysical characteristics, land use history; Fischer *et al*. 2008) and at regional scales, elements of both strategies will be needed to maintain connectivity among protected areas and to support the flow of organisms that provide ecosystem services to agricultural lands (Kremen 2015; Grass *et al*. 2019; Garibaldi *et al*. 2021). Identifying which landscapes may be best suited to each strategy therefore represents a spatial conservation challenge. For instance, areas where both agricultural productivity and connectivity are high may provide key opportunities for incentive programs that promote both food production and the flow of organisms through wildlife-friendly farming practices. Alternatively, landscapes with high potential for long-term food production but relatively limited connectivity value may be good candidates for government programs that keep lands in production and protect against conversion to other land uses (e.g., urbanization).

To explore the importance of agricultural lands in supporting connectivity across the United States, we modeled potential net movement of organisms across all terrestrial landscapes in CONUS using a circuit theory-based connectivity modeling approach (McRae *et al*. 2008; Dickson *et al*. 2019). We then used our connectivity results and existing information on agricultural land quality across CONUS to identify conservation opportunities on agricultural lands that balance species connectivity and long-term food security. To facilitate use of these results by landowners, conservation advocates, and government agencies, we developed an interactive web map, which allows users to explore the novel spatial data generated by our analysis and provides guidance on using these layers to identify conservation opportunities.

## Methods

To model connectivity across CONUS, we adopted an approach based on landscape structure, with less-modified landscapes assumed to support greater ecological flow (Dickson *et al*. 2017; Marrec *et al*. 2020). We tailored model parameters (e.g., maximum movement distances) to best reflect non-volant terrestrial vertebrates. Previous authors have noted that agricultural landscapes may represent ‘invisible mosaics’ (Fahrig *et al*. 2011), with a particular land cover category (e.g., cropland) actually representing a range of impacts on animal movement due to variation in management practices such as fertilizer application or cropping intensity. Here we build upon existing large-scale connectivity studies (e.g., McGuire *et al*. 2016; Dickson *et al*. 2017; Littlefield *et al*. 2017) by explicitly incorporating estimates of agricultural management intensity on cropland and pasture when determining landscape resistance to movement, using a novel method based on variation in vegetation cover during the growing season.

### Estimating human land use intensity

To evaluate the influence that agricultural lands and other modified landscapes exert on ecological flow, we estimated human land use intensity (*L*) for all locations (i.e., pixels in a gridded landscape) across CONUS. Our estimates of human land use intensity were based on a procedure originally described by Theobald (2013), which assigns literature-supported values of intensity to multiple forms of human land use and integrates these values into a single spatial data layer ranging from 0 (unmodified, ‘natural’) to 1 (heavily modified). Similar human land use intensity layers have formed the basis of previous ecological flow-based connectivity models (e.g., Dickson *et al*. 2017; Marrec *et al*. 2020).

To quantify land use intensity on agricultural lands, we started with existing, static *L* estimates for individual agricultural cover types (Theobald 2013) and incorporated a dynamic measure of management intensity based on temporal variation in vegetation cover at a given location. We used high spatial resolution (10 m) data on 2016 land cover from American Farmland Trust’s Farms Under Threat (FUT) analysis, which integrates data from multiple national-scale datasets to define several agricultural and non-agricultural cover classes (CSP 2020; data accessible from csp-fut.appspot.com). We focused on the four agricultural cover classes, which together account for 3.64 million km^2^, or approximately 47.6% of CONUS land area (Fig. 1). These agricultural classes are cropland (1,549,077 km^2^ across CONUS), pasture (430,369 km^2^), rangeland (1,658,472 km^2^), and woodland (174,323 km^2^). The woodland class is a subset of the Natural Resources Inventory forest class defined as “natural or planted forested cover that is part of a functioning farm unit” and is no more than 160 m from cropland or pasture (CSP 2020). We assigned each of these four classes with a baseline value of land-use intensity (*L*) corresponding with the general level of human disturbance associated with that agricultural type. For cropland and pasture, baseline *L* values of 0.5 and 0.4, respectively, were taken from Theobald (2013). Similar approaches to modeling ecological flow and/or landscape integrity have treated rangelands as having lower impact than cropland or pasture because rangelands tend to retain some natural vegetation cover and have relatively limited human influence (Buttrick *et al*. 2015; McRae *et al*. 2016). Woodlands are similarly characterized by relatively natural vegetation cover, albeit in close proximity to managed agricultural lands. We therefore assigned a baseline *L* value of 0.2 to both rangelands and woodlands in an effort to capture the greater potential for wildlife movement through these cover types.

**Figure 1.**
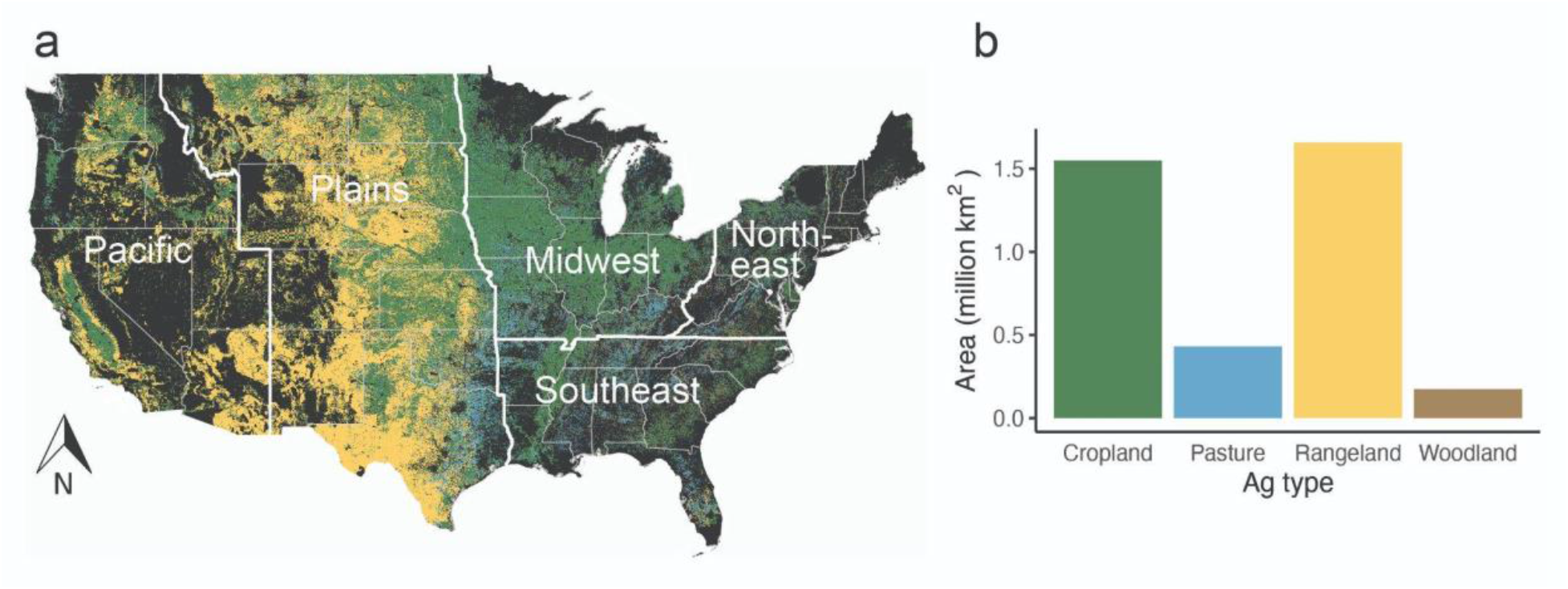
Agricultural land cover/use across the conterminous United States (CONUS). (**a**) Agricultural land cover is mapped across CONUS, with bold white lines and labels denoting Agricultural Research Service (ARS) regions. (**b**) Total area of each agriculture type across CONUS.

For both cropland and pasture, the agricultural cover types characterized by relatively intensive human management, we allowed *L* values to vary between pixels of the same cover type based on estimates of management intensity. Management intensity estimates were derived from temporal variability in vegetation cover based on the assumption that more intensively managed areas (e.g., croplands with high fertilizer inputs and/or multiple harvests per year; pasture subject to a high mowing frequency) will have greater variability in vegetation cover during the growing season than areas subject to less human intervention (e.g., fallow fields) (Franke *et al*. 2012; Gómez Giménez *et al*. 2017). We used a timeseries of Normalized Difference Vegetation Index (NDVI) values to estimate vegetation cover variability, acquiring cloud-free NDVI estimates at 16-day intervals from NASA’s MODIS system (MOD13Q1 products). For each cropland and pasture pixel across CONUS, we used NDVI estimates over a five year period (2014-2018) centered on 2016, the year of our land cover dataset. NDVI estimates were acquired during the growing season for each year, with growing season start and end dates defined separately for each U.S. state based on the planting dates database developed by Sacks et al. (2010) (see Appendix A for details).

For each pixel of cropland and pasture, we calculated the coefficient of variation for all NDVI values across the time series (hereafter, cvNDVI) as our estimate of vegetation cover variability. The coefficient of variation was chosen to account for differences between vegetation types (e.g., different crops) and geographic location in average plant greenness. For each cover type (cropland or pasture), we centered cvNDVI values by first calculating the mean for all pixels of that cover type within the same USDA plant hardiness zone (PHZ; (USDA 2012)) and then subtracting this mean value from the value for each pixel. PHZs describe bands of average annual minimum winter temperature across CONUS. We centered cvNDVI values based on means within a PHZ to account for potential differences in vegetation cover variability across latitudes and climatic conditions (e.g., lower variability in areas with shorter growing seasons). Averages (± SD) of mean-centered cvNDVI were -0.05 (± 0.13) and -0.03 (± 0.10) for cropland and pasture, respectively. To derive the final *L* value for cropland and pasture pixels, mean-centered cvNDVI values were added to the baseline *L* value for each cover type (0.5 for cropland and 0.4 for pastureland, see above), resulting in a range of final *L* estimates centered on the baseline value. Thus, pixels with lower than average vegetation cover variability for a given cover type and PHZ (i.e., negative mean-centered cvNDVI) received *L* values below the baseline value for that cover type and those with higher than average variability (positive mean-centered cvNDVI) received *L* values above the baseline. For rangeland and woodland, we did not incorporate vegetation index data into *L* estimates, instead using baseline *L* values for all pixels under the assumption that variability in NDVI will be more strongly associated with phenology and plant community composition than with human management intensity in the cover types characterized by relatively natural vegeation.

We tested the validity of cvNDVI as a proxy for agricultural management intensity by comparing cvNDVI values between agricultural cover types; between irrigated, unirrigated, and fallow cropland; and across a gradient of nitrogen fertilizer use. These validation analyses are described in Appendix B. The validation steps confirmed the utility of cvNDVI as a proxy for management intensity, showing that (i) cropland pixels had significantly higher average cvNDVI than pasture; (ii) for both cropland and pasture, cvNDVI was positively correlated with nitrogen fertilizer usage; and (iii) among cropland pixels, irrigated crops had the highest average cvNDVI, followed by unirrigated crops and then fallow fields (Appendix B).

To create a comprehensive layer of human land use intensity across CONUS, we combined our novel agriculture *L* layer with layers describing other forms of human land use, and incorporated the impact of nearby land uses and disturbances on a given location by allowing the value of each pixel to extend beyond the focal pixel itself. For all non-agricultural land uses we used an existing *L* model (CSP 2019) that integrates multiple land use variables into three human impact categories - urban (including data on residential development and nighttime lights), transportation (including roads, railways, powerlines, and pipelines), and energy (including oil and gas wells, coal mines, and utility-scale solar and wind installations). Details on the development of the final *L* layer (*L*_*all*_) are provided in Appendix A.

### Estimating landscape resistance and modeling connectivity

We used our final land use intensity layer, *L*_*all*_, to derive a landscape resistance surface, which estimates the difficulty an organism experiences in moving through each pixel on the landscape (Zeller *et al*. 2012). Following Dickson et al. (2017), who conducted a sensitivity analysis to determine an appropriate formula for deriving resistance surfaces by rescaling *L* values, we calculated resistance (*R*) as

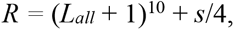

where *s* is the percent slope of a given pixel, thus penalizing areas of steep slope to account for the effects of substantial elevational changes on the movement capacity of many terrestrial species (Dickson *et al*. 2017). This resulted in resistance values ranging between 1 (natural landscape) and 1032 (heavily modified landscape). We assigned all water bodies greater than approximately 100 m across a resistance value of 1000 to reflect the difficulty of moving through water for terrestrial species. The above rescaling formula led to a relatively high contrast between the resistance values assigned to locations with low, medium, and high *L*_*all*_ values. For comparison, we derived a second resistance surface using a low-contrast rescaling formula suggested by Marrec et al. (2020). See Appendix C for a comparison of the two resistance surfaces and resulting connectivity models.

We modeled source strength, i.e., the predicted probability or intensity of movement from a given location on the landscape (McRae *et al*. 2008, 2016), as the degree of ecological intactness of a given pixel, which we calculated as 1 - *L*_*all*_. Our source strength layer therefore ranged between 0 and 1, with relatively intact habitats receiving values close to 1, while partially modified landscapes (e.g., agricultural lands) received low but non-zero values. We assigned zero source strength to areas unlikely to represent sources of terrestrial animal movement, using the 2016 National Land Cover Data Base (NLCD; Dewitz 2019) to set pixels categorized as developed, open water, perennial snow/ice, or barren rock (i.e., all NLCD cover classes < 40) to zero. Resistance and source strength rasters for CONUS were derived at 250-m resolution using Google Earth Engine (GEE; Gorelick *et al*. 2017).

Following McRae et al. (2016), we ran omni-directional connectivity models across CONUS using the Omniscape algorithm. Omniscape is based on concepts from electronic circuit theory (McRae *et al*. 2008; Dickson *et al*. 2019), modeling the movement of organisms across the landscape as the flow of electrical current through a circuit. Omniscape allows users to fit “coreless” connectivity models in which every pixel may potentially serve as a source and/or target of movement, rather than only modeling connectivity between habitat cores, and thus allowing current to potentially flow in all directions. The algorithm uses a moving window approach, iteratively treating every pixel in the source strength layer with a value greater than zero as a target for electrical current and connecting that pixel to all other non-zero pixels within the moving window radius, which serve as current sources. Current is then injected into the source pixels (with the amount of current proportional to source strength) and flows across the resistance surface (McRae *et al*. 2016; Landau *et al*. 2021). The cumulative current flow across all iterations of the moving window provides an estimate of the probability or intensity of the movement of organisms through every pixel on the landscape. The moving window radius is a key parameter, setting the maximum movement distance (i.e., the maximum distance between source and target pixels). Here we used a radius of 150 km, which approximates the upper dispersal distances of many large-bodied terrestrial vertebrates (Sutherland *et al*. 2000). To increase processing speed, we only treated every forty-first pixel as a (potential) target in the moving window. For comparison, we also ran connectivity models using smaller moving window radii, as described in Appendix C. Connectivity models were run in the Omniscape.jl software package in Julia (Landau et al. 2021).

We summarized cumulative current flow values from the Omniscape model within regions of the U.S. (defined by the USDA Agricultural Research Service [ARS]), and compared current flow on agricultural lands with that on other land cover/land use types (including developed and natural lands), providing an overview of agricultural land contributions to connectivity across the country. These analyses are described in detail in Appendix A. To further explore the drivers of high or low connectivity values on agricultural lands, we also estimate the total amount of natural land cover and development (based on NLCD categories) within a 1-km radius of each location on agricultural lands, hypothesizing that agriculture surrounded by greater amounts of natural land cover and lower levels of development would tend to have higher current flow. We tested the effect of surrounding land cover/land use on agricultural land current flow using a spatial error regression analysis (Dale and Fortin 2014) described in Appendix A.

### Identifying conservation opportunities on agricultural lands

To help identify and prioritize conservation opportunities on agricultural lands across CONUS, we categorized all agricultural pixels based on both their potential to support ecological flow and their value for long-term food production. This analysis drew upon results of the connectivity model described above as well as a CONUS-wide data layer estimating productivity, versatility, and resilience (PVR; 10-m resolution) of agricultural lands circa 2016, which was developed as part of the FUT analysis described above (CSP 2020; viewable at csp-fut.appspot.com). PVR quantifies the long-term sustainability of maintaining a given area in cultivation based on soil and land cover characteristics and the type of agriculture practiced at a given location. See Appendix A for further details.

We masked the current flow and PVR datasets to only agricultural pixels and calculated quantiles of each dataset to identify pixels falling into ‘low’ (< 33% quantile), ‘medium’ (33% to 66%), and ‘high’ (> 66%) categories for connectivity and PVR. Because some regions have generally higher connectivity or PVR values than others, calculation of quantiles was conducted separately for each ARS region across CONUS. Thus, both connectivity and PVR values are considered relative to other agricultural pixels in the same region. We used these quantile estimates to generate a pixel-level bivariate map in which each pixel on agricultural land was ranked as ‘low’, ‘medium’, or ‘high’ for both connectivity and PVR. We used this map to link joint connectivity and food production value with specific conservation opportunities and/or financial incentives administered by the USDA.

## Results

The importance of agricultural lands in facilitating the movement of organisms varied substantially across regions of the U.S. (Fig. 2, Table 1). At the ARS regional level, intensively cropped landscapes in the midwest (e.g., southern Minnesota, Iowa, and Illinois, Fig. 2) exhibited relatively high resistance to movement (Appendix D, Fig. D1) and low connectivity (Fig. 2, Table 1), while regions with extensive rangelands (e.g., central Nebraska, southwestern Texas) exhibited lower landscape resistance and were characterized by diffuse but relatively high current flow, comparable to natural landscapes in the western U.S. (e.g., central Nevada, northern Idaho; Fig. 2, Table 1). The plains region had the largest proportion of total land area in agriculture (70.0%) and, consequently, the greatest contribution of agricultural lands to overall connectivity in the region - agricultural lands accounted for 17.6% of top connectivity lands (i.e., lands in the top quartile of current flow, Appendix A) in the plains region (Table 1).

**Table 1.**
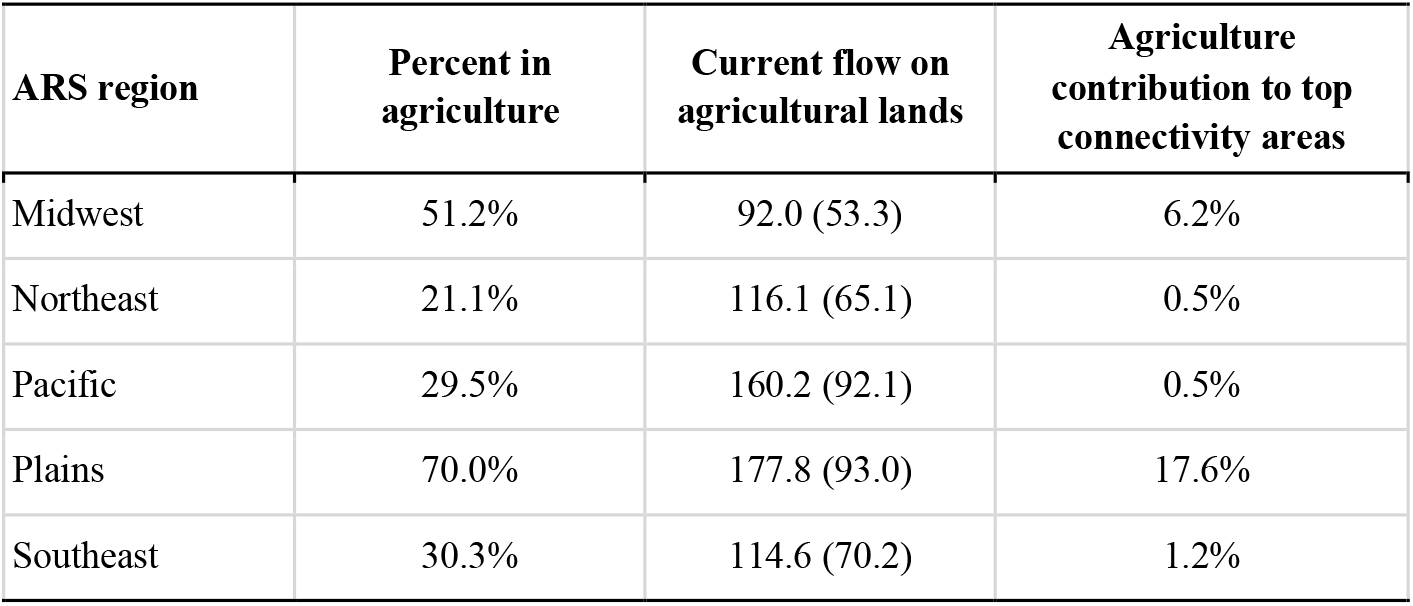
Summary of connectivity on agricultural lands by Agricultural Research Service (ARS) region. *Percent in agriculture*: amount of total land area in the region categorized as cropland, pasture, rangeland, or woodland. *Current flow on agricultural lands*: mean (standard deviation) of current flow across all agricultural pixels in the region. *Agriculture contribution to top connectivity areas*: proportion of top lands for connectivity in the region (i.e., those with current flow values falling within the top quartile for the region) occurring on agricultural lands. ARS regions are shown in Fig. 1a.

| ARS region | Percent in agriculture | Current flow on agricultural lands | Agriculture contribution to top connectivity areas |
| --- | --- | --- | --- |
| Midwest | 51.2% | 92.0 (53.3) | 6.2% |
| Northeast | 21.1% | 116.1 (65.1) | 0.5% |
| Pacific | 29.5% | 160.2 (92.1) | 0.5% |
| Plains | 70.0% | 177.8 (93.0) | 17.6% |
| Southeast | 30.3% | 114.6 (70.2) | 1.2% |

**Figure 2.**
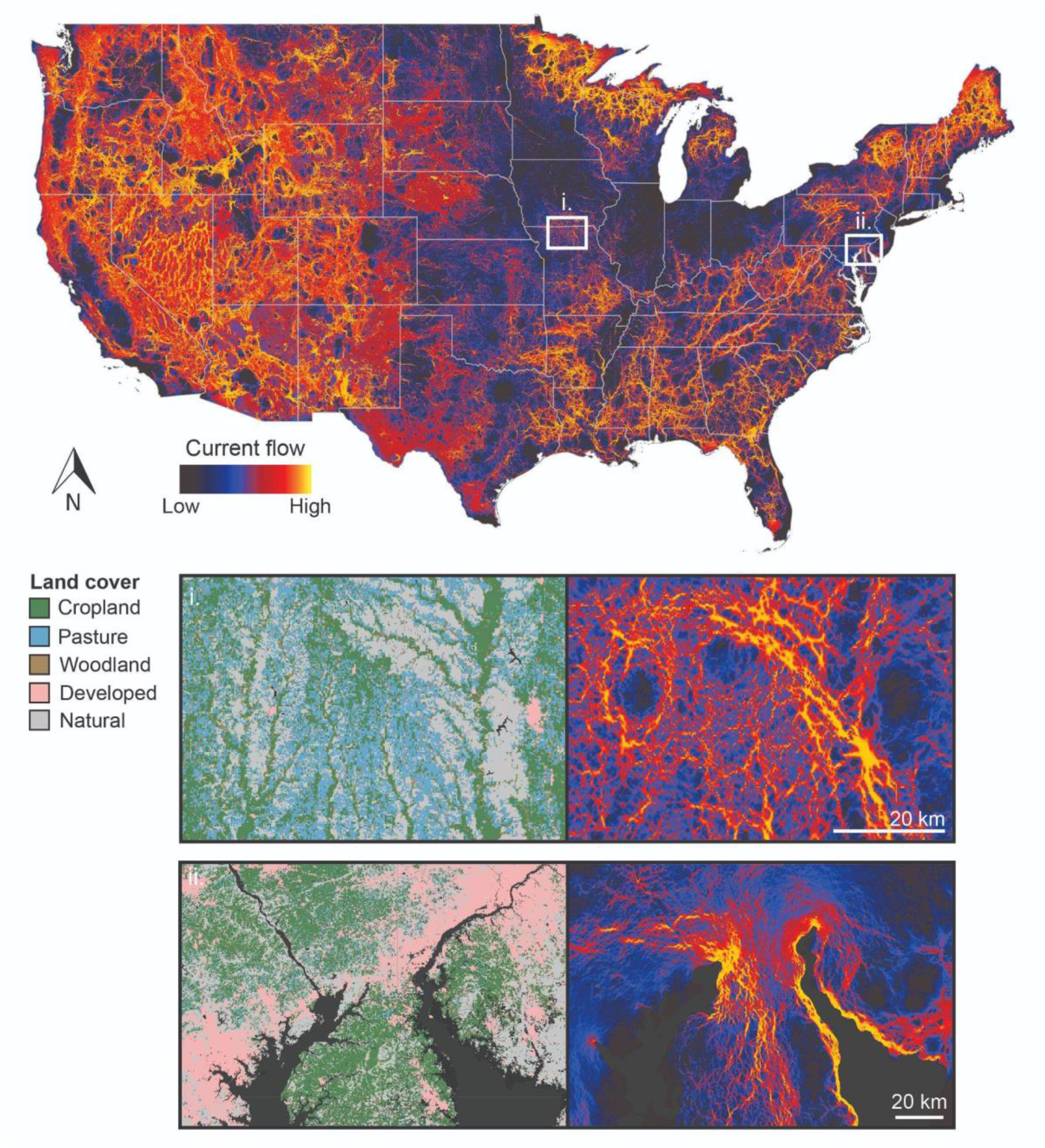
Map of connectivity (current flow) across the conterminous United States. Insets show details of agricultural landscapes with high connectivity value in (i) the central Midwest (northern Missouri) and (ii) the coastal Northeast (Delmarva peninsula, near the border of Maryland, Delaware, and Pennsylvania), where the persistence of patches of natural vegetation (forest fragments and strips of riparian or coastal vegetation) positively influence the current flow values of neighboring agricultural lands.

Current flow on agricultural lands tended to be intermediate between current flow values of more developed landscapes (e.g., urban and suburban areas) and those of natural areas (including GAP 1 and 2 protected areas; Fig. 3a). The amount of natural and developed land in the vicinity of agricultural lands substantially influenced the connectivity value of individual agricultural pixels. Our top spatial error regression model (Table D1, ΔAIC of next best model = 114.3) sufficiently accounted for spatial autocorrelation in model residuals (Moran’s I = -0.03, *p* = 0.99) and included an interaction between agricultural cover type and the non-linear effects of surrounding land cover/use. Current flow values on all agriculture types were positively influenced by the amount of natural vegetation within 1 km (Fig. 3b, see also Fig. 2 insets) and negatively influenced by the amount of developed land within 1 km (Fig. 3c).

**Figure 3.**
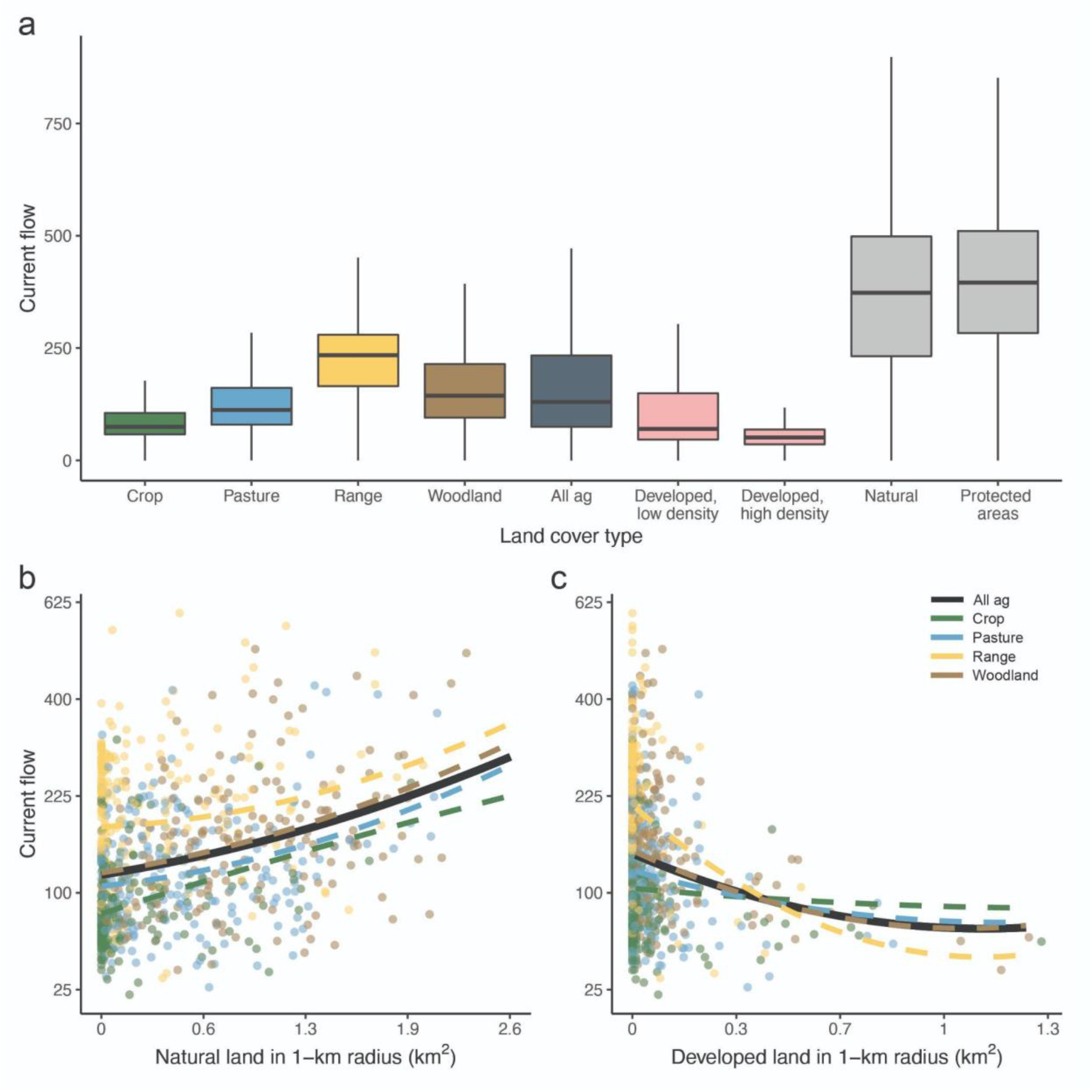
Current flow across land cover/use categories. (**a**) The range of current flow values on agricultural lands (cropland, pasture, rangeland, woodland, and all agricultural categories combined [‘all ag’]) is compared to that of developed areas and landscapes characterized by more natural land cover (i.e., all natural lands and lands within USGS GAP 1 and GAP 2 protected areas). Data are summarized as standard boxplots with whiskers representing 1.5 times the interquartile range. Outliers are excluded for clarity (see Appendix D, Figure D2 for a version with all outliers shown). Current flow values on agricultural land cover types are influenced by surrounding land cover/use, including the amount of (**b**) natural lands and (**c**) developed lands within 1 km. Fitted lines in **b** and **c** show the mean relationship between surrounding land cover and current flow as estimated by a spatial error model. For clarity, a randomly selected subset (n = 800) of data points used in the analysis are shown.

Mapping the combined rankings of current flow and PVR (Fig. 4) revealed that 2.7% of all agricultural lands (10.2 million hectares [Mha]) have high values for both connectivity and PVR (i.e., within the top 33% of agricultural lands in the same region; Table 2). The proportion of lands in this ‘high-high’ category varied between ARS regions, being lowest in the Plains (1.3%) and highest in the Northeast (5.8%, Table 2, Fig. 4 inset). Areas of low connectivity (i.e., values in the bottom 33%) and high PVR were more common overall, accounting for 21.3% of all agricultural lands across CONUS (81.3 Mha), followed by areas of high connectivity and low PVR (15.5%, 59.3 Mha) and areas in the lowest category for both connectivity and PVR (4.2%, 16.2 Mha; Table 2).

**Table 2.**
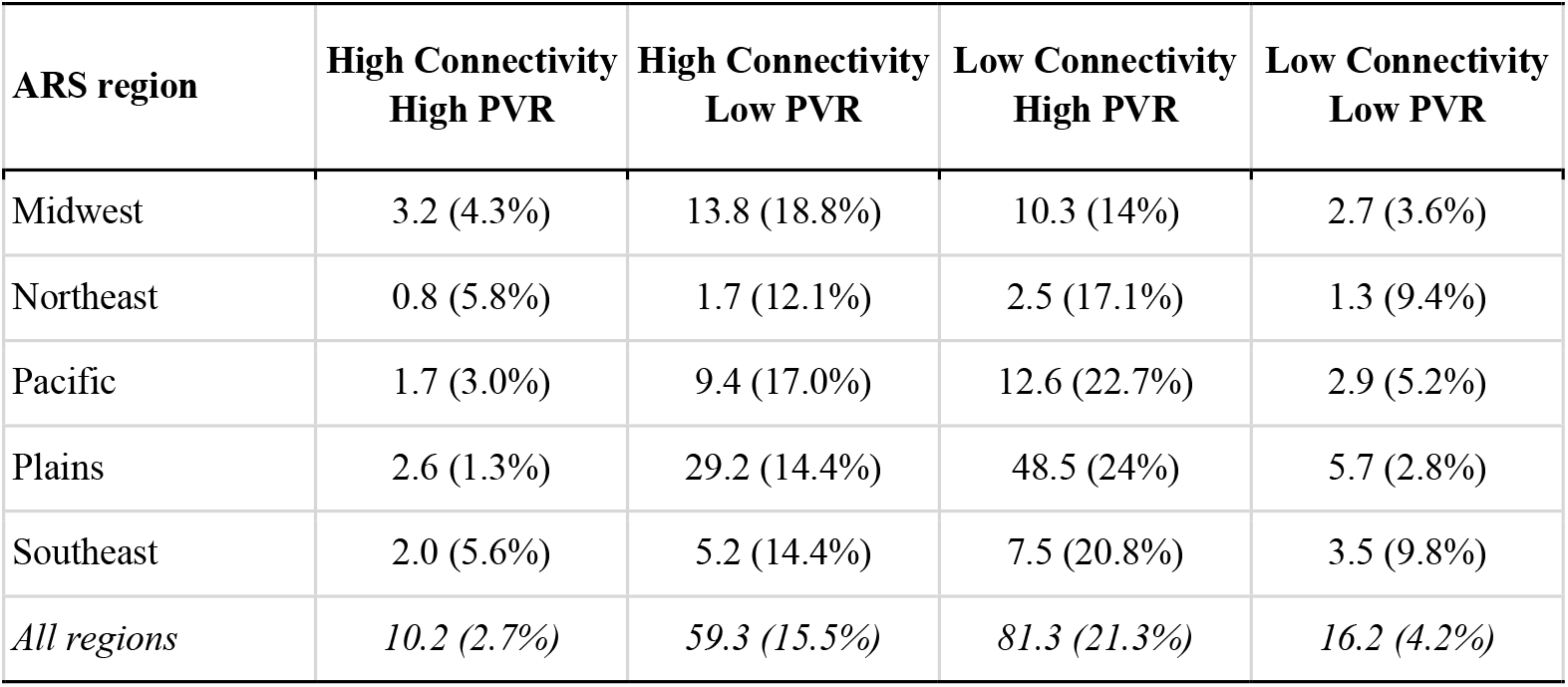
Total area (Mha) and percentages of agricultural lands in each Agricultural Research Service (ARS) region falling into high (i.e., values within the top 33% of all agricultural lands in the same region) and low (values in the bottom 33%) rankings for connectivity and PVR. Values are also given for all regions combined (i.e., all agricultural lands across the conterminous United States). ARS regions are shown in Fig. 1a.

| ARS region | High Connectivity<br>High PVR | High Connectivity<br>Low PVR | Low Connectivity<br>High PVR | Low Connectivity<br>Low PVR |
| --- | --- | --- | --- | --- |
| Midwest | 3.2 (4.3%) | 13.8 (18.8%) | 10.3 (14%) | 2.7 (3.6%) |
| Northeast | 0.8 (5.8%) | 1.7 (12.1%) | 2.5 (17.1%) | 1.3 (9.4%) |
| Pacific | 1.7 (3.0%) | 9.4 (17.0%) | 12.6 (22.7%) | 2.9 (5.2%) |
| Plains | 2.6 (1.3%) | 29.2 (14.4%) | 48.5 (24%) | 5.7 (2.8%) |
| Southeast | 2.0 (5.6%) | 5.2 (14.4%) | 7.5 (20.8%) | 3.5 (9.8%) |
| *All regions* | *10.2 (2.7%)* | *59.3 (15.5%)* | *81.3 (21.3%)* | *16.2 (4.2%)* |

**Figure 4.**
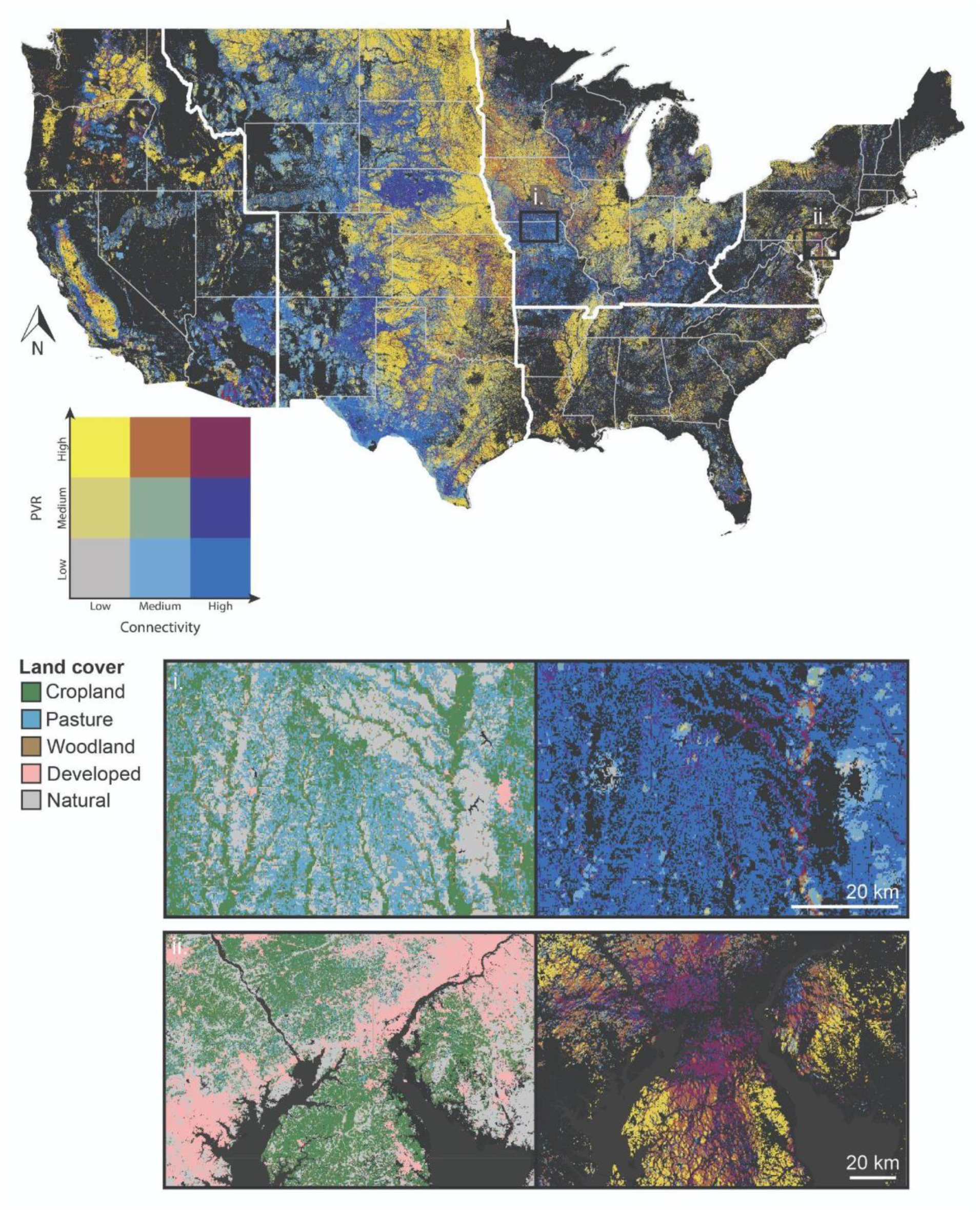
Map of agricultural lands ranked based on quantiles of connectivity (i.e., current flow) and productivity, versatility, and resilience (PVR), a measure of agricultural land quality. White lines indicate the boundaries of Agricultural Research Services (ARS) regions, within which quantiles of connectivity and PVR were calculated (color scales are therefore relative to other pixels in the same ARS region). Non-agricultural land cover/use types are shown in black. The insets show details of agricultural landscapes with high connectivity value and either high or low PVR, and correspond to those shown in Figure 2. The central Midwest landscape (i) is characterized by relatively high connectivity value but low PVR (deep blue color). The northern end of the Delmarva Peninsula (ii) has large amounts of land with high connectivity and high PVR (maroon color).

## Discussion

Our results highlight the potential for agricultural lands across the U.S. to provide important movement routes for terrestrial species, supporting connectivity through otherwise heavily modified landscapes. Current flow through agricultural pixels depended strongly on the type of agriculture practiced at a given location and the intensity of human land use in the surrounding landscape. Generally, agricultural lands supported greater current flow than developed areas, suggesting an important role for agricultural lands as corridors linking areas of high-quality habitat. As the human footprint continues to expand, moderately impacted landscapes such as agricultural fields and grazing lands will be increasingly important movement habitat for many species (Suraci *et al*. 2020). Identifying conservation opportunities on agricultural lands is therefore critical to preventing further biodiversity loss while promoting long-term food security (Kremen and Merenlender 2018; Leclère *et al*. 2020).

Our maps reveal that landscapes ranking in the highest quantile for both connectivity and PVR constitute 10.2 Mha (approximately 3%) of agricultural lands across CONUS. These areas represent key opportunities for ‘land-sharing’ programs that promote biodiversity and food production on the same land holdings (Fischer *et al*. 2008, 2014; Kremen 2015; Garibaldi *et al*. 2021) by incentivizing wildlife-friendly farming practices (e.g., USDA’s Conservation Stewardship Program and Environmental Quality Incentives Program [EQIP]) and protecting agricultural lands against conversion to more intensive land uses (e.g., USDA’s Agricultural Conservation Easement Program [ACEP]). Increasing the capacity for farmland in this ‘high connectivity-high PVR’ category to support native species and wildlife movement - for instance, by planting non-crop vegetation strips along field edges and increasing crop diversity (Kremen *et al*. 2012) - may be critical for preserving connectivity in some areas of the U.S. and is consistent with recent proposals to increase conservation efforts on private lands (e.g., the Biden administration’s commitment to conserve 30% of U.S. lands by 2030; Exec. Order No. 14008, 2021). Importantly, recent work has shown that incorporating such wildlife-friendly farming practices can stabilize (Gaudin *et al*. 2015) or even increase agricultural yields (Pywell *et al*. 2015), setting up the potential for biodiversity and food production ‘win-wins’ (Mitchell *et al*. 2013).

Our model results also identified substantial amounts of agricultural land across CONUS that are of high value for either connectivity (i.e., high connectivity-low PVR; 59.3 Mha in total) or food production (i.e., low connectivity-high PVR; 81.3 Mha) but not both. Such landscapes are particularly common in the Plains region, where large expanses of land with high PVR are devoted to intensive crop production (thus limiting connectivity value), but are interspersed with areas of less intensive agriculture on lower productivity lands. Such landscapes may be excellent candidates for a combination of management policies that reflect a ‘land-sparing’ conservation strategy (Phalan *et al*. 2011; Grass *et al*. 2019). Lands in the ‘low connectivity-high PVR’ category could be targeted for programs that keep lands in production and protect against conversion to development (e.g., ACEP), thus safeguarding the most productive agricultural lands. Meanwhile, neighboring areas in the ‘high connectivity-low PVR’ category could be maintained as low-intensity agriculture (pasture or rangelands) through enrollment in programs that support grazing (e.g., EQIP, CSP or USDA’s term-limited Grasslands Conservation Reserve Program [CRP]) and/or through permanent easements (e.g., via ACEP). Where appropriate, such areas could also be considered for removal from production in favor of ecological restoration to support habitat and movement of native species (e.g., through USDA’s Conservation Reserve Enhancement Program). Under such a conservation strategy, high-connectivity lands maintained as low-intensity agriculture or removed from agriculture altogether can act as connectivity ‘stepping stones’ (Wimberly *et al*. 2018; Doherty and Driscoll 2018) to support species movement through otherwise intensively managed landscapes and connect larger patches of high-quality habitat (e.g., protected areas). It is critical that any such land-sparing strategy be implemented at a relatively large spatial scale (i.e., across multiple land holdings within a region) to ensure sufficient connectivity across a network of ‘spared’ habitat patches to support dispersal and patch colonization and to prevent isolation of the larger protected areas that such habitat patches connect (Lamb *et al*. 2016; Grass *et al*. 2019).

Spatial context plays a substantial role in determining the connectivity value of agricultural lands. Our spatial regression analysis showed that, regardless of agriculture type, current flow on agricultural pixels was highest when those pixels were embedded in a broader landscape consisting of large amounts of natural land cover. This finding is consistent with previous work showing that biodiversity in agricultural systems tends to be higher in heterogeneous landscapes consisting of a mix of agricultural and non-agricultural cover types (e.g., crop fields and pastures interspersed with woodlots and riparian buffers) (Donald and Evans 2006; Fahrig *et al*. 2011; Reynolds *et al*. 2018; Kremen and Merenlender 2018). By promoting connectivity, the presence of natural land cover in agricultural systems can also directly benefit food production, providing ecosystem services such as pollination and biological pest control through the (re)colonization and spillover of service-providing organisms from natural to cultivated patches (Blitzer *et al*. 2012; Kormann *et al*. 2016; Grass *et al*. 2019). Therefore, maintaining or restoring natural vegetation within agricultural systems is likely to have benefits across scales, promoting biodiversity and the provisioning of ecosystem services at the local level of individual farms while facilitating regional-scale connectivity across networks of working lands and protected areas.

Structural connectivity models such as the one used here are typically aimed at describing connectivity for a wide range of species (Dickson *et al*. 2017; Marrec *et al*. 2020) and perform well in terms of their overlap with focal species connectivity models, particularly for larger-bodied species or those with high movement capacity (Krosby *et al*. 2015). However, it is important to note that our model was not calibrated to the movement or habitat preferences of any particular focal species and thus may not fully capture the best movement pathways for a given species of interest. An important next step for connectivity conservation on agricultural lands will be to adapt the methods developed here in building connectivity models for focal species of conservation concern. Our NDVI-based approach to capturing variation in management intensity within a given agricultural cover type could readily be adapted to focal species functional connectivity models. Researchers can use species distribution models (Keeley *et al*. 2016) or resource selection functions (Zeller *et al*. 2014) to quantify the effect of agricultural management intensity on species habitat suitability or probability of use and translate these values into landscape resistance (Zeller *et al*. 2012; Suraci *et al*. 2020). Such efforts will be important for species-level management and will likely reveal substantial diversity in the capacity of agricultural lands to provide habitat value for individual species (Phalan *et al*. 2011; Reynolds *et al*. 2018).

We expect that our results will be useful in prioritizing conservation actions across a range of scales. At the local level, farmers, land trusts, and conservation advocates can use information on the joint value of agricultural lands for connectivity and food production to identify site-specific conservation strategies (e.g., land sparing vs. sharing), explore the types of conservation-focused financial incentives applicable to the landscapes they work in, and advocate for policies that support the conservation of working lands. At the state and federal levels, agencies tasked with administering agricultural conservation programs can use these results to better target funding to areas likely to have the greatest impact for promoting biodiversity and food security, ideally employing a landscape-scale approach that leads to heterogeneous agricultural-natural mosaics that benefit both producers and native species (Kremen *et al*. 2012). To help facilitate planning and conservation action based on our results, we have developed an interactive web application (https://cspbeta.z22.web.core.windows.net/), allowing users to visualize the spatial data developed here within their regions of interest. We hope that these tools can contribute to a collaborative process between landowners, governments, and conservationists to design landscapes that support both native species and a sustainable food supply.

## Author contributions

JPS, BDG, MCH, and AS conceived of the study. JPS led the analysis and writing with substantial help from CEL and CCN. All authors provided feedback on initial drafts.

## Acknowledgements

We thank V Landau, J Anderson, K Carroll, and T Nogeire-McRae for their helpful feedback on this work.

## References

Blitzer EJ, Dormann CF, Holzschuh A, et al. 2012. Spillover of functionally important organisms between managed and natural habitats. Agriculture, Ecosystems & Environment 146: 34–43.

Buttrick S, Popper K, Schindel M, et al. 2015. Conserving Nature’s Stage: Identifying Resilient Terrestrial Landscapes in the Pacific Northwest. The Nature Conservancy, Portland, Oregon.

CSP. 2019. Methods and approach used to estimate the loss and fragmentation of natural lands in the conterminous U.S. from 2001 to 2017. Truckee, CA.

CSP. 2020. Description of the approach, data, and analytical methods used for the Farms Under Threat: State of the States project, version 2.0. Final Technical Report. Truckee, CA.

Dale MRT and Fortin M-J. 2014. Spatial Analysis: A Guide For Ecologists. Cambridge University Press.

Dewitz J. 2019. National Land Cover Database (NLCD) 2016 Products: U.S. Geological Survey data release.

Dickson BG, Albano CM, Anantharaman R, et al. 2019. Circuit-theory applications to connectivity science and conservation. Conservation Biology 33: 239–49.

Dickson BG, Albano CM, McRae BH, et al. 2017. Informing strategic efforts to expand and connect protected areas using a model of ecological flow, with application to the western United States. Conservation Letters 10: 564–71.

Doherty TS and Driscoll DA. 2018. Coupling movement and landscape ecology for animal conservation in production landscapes. Proceedings of the Royal Society B: Biological Sciences 285: 20172272.

Donald PF and Evans AD. 2006. Habitat connectivity and matrix restoration: the wider implications of agri-environment schemes. Journal of Applied Ecology 43: 209–18.

Fahrig L, Baudry J, Brotons L, et al. 2011. Functional landscape heterogeneity and animal biodiversity in agricultural landscapes. Ecology Letters 14: 101–12.

Fischer J, Abson DJ, Butsic V, et al. 2014. Land Sparing Versus Land Sharing: Moving Forward. Conservation Letters 7: 149–57.

Fischer J, Brosi B, Daily GC, et al. 2008. Should Agricultural Policies Encourage Land Sparing or Wildlife-Friendly Farming? Frontiers in Ecology and the Environment 6: 380–5.

Franke J, Keuck V, and Siegert F. 2012. Assessment of grassland use intensity by remote sensing to support conservation schemes. Journal for Nature Conservation 20: 125–34.

Garibaldi LA, Oddi FJ, Miguez FE, et al. 2021. Working landscapes need at least 20% native habitat. Conservation Letters 14: e12773.

Gaudin ACM, Tolhurst TN, Ker AP, et al. 2015. Increasing Crop Diversity Mitigates Weather Variations and Improves Yield Stability. PLOS ONE 10: e0113261.

Gómez Giménez M, Jong R de, Della Peruta R, et al. 2017. Determination of grassland use intensity based on multi-temporal remote sensing data and ecological indicators. Remote Sensing of Environment 198: 126–39.

Gorelick N, Hancher M, Dixon M, et al. 2017. Google Earth Engine: Planetary-scale geospatial analysis for everyone. Remote Sensing of Environment 202: 18–27.

Grass I, Loos J, Baensch S, et al. 2019. Land-sharing/-sparing connectivity landscapes for ecosystem services and biodiversity conservation. People and Nature 1: 262–72.

Heller NE and Zavaleta ES. 2009. Biodiversity management in the face of climate change: A review of 22 years of recommendations. Biological Conservation 142: 14–32.

Hunter MC, Smith RG, Schipanski ME, et al. 2017. Agriculture in 2050: Recalibrating Targets for Sustainable Intensification. BioScience 67: 386–91.

Keeley ATH, Beier P, and Gagnon JW. 2016. Estimating landscape resistance from habitat suitability: effects of data source and nonlinearities. Landscape Ecol 31: 2151–62.

Kormann U, Scherber C, Tscharntke T, et al. 2016. Corridors restore animal-mediated pollination in fragmented tropical forest landscapes. Proceedings of the Royal Society B: Biological Sciences 283: 20152347.

Kremen C. 2015. Reframing the land-sparing/land-sharing debate for biodiversity conservation. Annals of the New York Academy of Sciences 1355: 52–76.

Kremen C, Iles A, and Bacon C. 2012. Diversified Farming Systems: An Agroecological, Systems-based Alternative to Modern Industrial Agriculture. Ecology and Society 17.

Kremen C and Merenlender AM. 2018. Landscapes that work for biodiversity and people. Science 362.

Krosby M, Breckheimer I, John Pierce D, et al. 2015. Focal species and landscape “naturalness” corridor models offer complementary approaches for connectivity conservation planning. Landscape Ecol 30: 2121–32.

Kummu M, Heino M, Taka M, et al. 2021. Climate change risks pushing one-third of global food production outside the safe climatic space. One Earth 4: 720–9.

Lamb A, Balmford A, Green RE, and Phalan B. 2016. To what extent could edge effects and habitat fragmentation diminish the potential benefits of land sparing? Biological Conservation 195: 264–71.

Landau VA, Shah VB, Anantharaman R, and Hall KR. 2021. Omniscape.jl: Software to compute omnidirectional landscape connectivity. Journal of Open Source Software 6: 2829.

Lark TJ, Spawn SA, Bougie M, and Gibbs HK. 2020. Cropland expansion in the United States produces marginal yields at high costs to wildlife. Nature Communications 11: 4295.

Leclère D, Obersteiner M, Barrett M, et al. 2020. Bending the curve of terrestrial biodiversity needs an integrated strategy. Nature 585: 551–6.

Littlefield CE, Krosby M, Michalak JL, and Lawler JJ. 2019. Connectivity for species on the move: supporting climate-driven range shifts. Frontiers in Ecology and the Environment 17: 270–8.

Littlefield CE, McRae BH, Michalak JL, et al. 2017. Connecting today’s climates to future climate analogs to facilitate movement of species under climate change. Conservation Biology 31: 1397–408.

Maas B, Brandl M, Hussain RI, et al. 2021. Functional traits driving pollinator and predator responses to newly established grassland strips in agricultural landscapes. Journal of Applied Ecology 58: 1728–37.

Marrec R, Abdel Moniem HE, Iravani M, et al. 2020. Conceptual framework and uncertainty analysis for large-scale, species-agnostic modelling of landscape connectivity across Alberta, Canada. Scientific Reports 10: 6798.

McGuire JL, Lawler JJ, McRae BH, et al. 2016. Achieving climate connectivity in a fragmented landscape. PNAS 113: 7195–200.

McRae BH and Beier P. 2007. Circuit theory predicts gene flow in plant and animal populations. PNAS 104: 19885–90.

McRae BH, Dickson BG, Keitt TH, and Shah VB. 2008. Using circuit theory to model connectivity in ecology, evolution, and conservation. Ecology 89: 2712–24.

McRae B, Popper K, Jones A, et al. 2016. Conserving nature’s stage: Mapping omnidirectional connectivity for resilient terrestrial landscapes in the Pacific Northwest. The Nature Conservancy, Portland, OR.

Mitchell MGE, Bennett EM, and Gonzalez A. 2013. Linking Landscape Connectivity and Ecosystem Service Provision: Current Knowledge and Research Gaps. Ecosystems 16: 894–908.

Newbold T, Hudson LN, Hill SLL, et al. 2015. Global effects of land use on local terrestrial biodiversity. Nature 520: 45–50.

Phalan B, Onial M, Balmford A, and Green RE. 2011. Reconciling Food Production and Biodiversity Conservation: Land Sharing and Land Sparing Compared. Science 333: 1289–91.

Polasky S, Nelson E, Camm J, et al. 2008. Where to put things? Spatial land management to sustain biodiversity and economic returns. Biological Conservation 141: 1505–24.

Pywell RF, Heard MS, Woodcock BA, et al. 2015. Wildlife-friendly farming increases crop yield: evidence for ecological intensification. Proc Biol Sci 282: 20151740.

Reynolds C, Fletcher RJ, Carneiro CM, et al. 2018. Inconsistent effects of landscape heterogeneity and land-use on animal diversity in an agricultural mosaic: a multi-scale and multi-taxon investigation. Landscape Ecol 33: 241–55.

Sacks WJ, Deryng D, Foley JA, and Ramankutty N. 2010. Crop planting dates: an analysis of global patterns. Global Ecology and Biogeography 19: 607–20.

Sohl TL, Sayler KL, Bouchard MA, et al. 2014. Spatially explicit modeling of 1992–2100 land cover and forest stand age for the conterminous United States. Ecological Applications 24: 1015–36.

Stanton RL, Morrissey CA, and Clark RG. 2018. Analysis of trends and agricultural drivers of farmland bird declines in North America: A review. Agriculture, Ecosystems & Environment 254: 244–54.

Suraci JP, Nickel BA, and Wilmers CC. 2020. Fine-scale movement decisions by a large carnivore inform conservation planning in human-dominated landscapes. Landscape Ecol 35: 1635–49.

Sutherland GD, Harestad AS, Price K, and Lertzman KP. 2000. Scaling of Natal Dispersal Distances in Terrestrial Birds and Mammals. Conservation Ecology 4.

Theobald DM. 2013. A general model to quantify ecological integrity for landscape assessments and US application. Landscape Ecol 28: 1859–74.

Tilman D, Clark M, Williams DR, et al. 2017. Future threats to biodiversity and pathways to their prevention. Nature 546: 73–81.

Tu Y, Chen B, Yu L, et al. 2021. How does urban expansion interact with cropland loss? A comparison of 14 Chinese cities from 1980 to 2015. Landscape Ecol 36: 243–63.

USDA. 2012. Plant Hardiness Zone Map. https://planthardiness.ars.usda.gov/.

Wimberly MC, Narem DM, Bauman PJ, et al. 2018. Grassland connectivity in fragmented agricultural landscapes of the north-central United States. Biological Conservation 217: 121–30.

Zeller KA, McGarigal K, Beier P, et al. 2014. Sensitivity of landscape resistance estimates based on point selection functions to scale and behavioral state: pumas as a case study. Landscape Ecol 29: 541–57.

Zeller KA, McGarigal K, and Whiteley AR. 2012. Estimating landscape resistance to movement: a review. Landscape Ecol 27: 777–97.

