## Appendix for "Mapping connectivity and conservation opportunity on agricultural lands across the conterminous United States"

**Contents**

*Appendix A.* Supplementary methods
*Appendix B.* Validating the relationship between vegetation index variability and management intensity
*Appendix C.* Sensitivity of connectivity model results to resistance surface scaling and moving window size.
*Appendix D.* Supplementary tables and figures
*Supplementary References*

**Appendix A. Supplementary methods**

*Defining growing season start and end dates for bounding NDVI time series data*

For each U.S. state, we used the planting dates database developed by Sacks et al. (2010) to extract the planting start date and harvest end date for a common crop type and used this as the date range within which NDVI estimates were acquired for all cropland and pasture pixels in that state. For consistency, we used corn as the crop type to define growing season dates for all states where corn is grown. For states where corn is not grown, we used another spring-planted crop represented in the database (spring barley in Nevada and potatoes in Maine, Massachusetts, and Rhode Island). For states that are absent from the database (Connecticut, Vermont, and New Hampshire), we used the planting start and harvest end dates for nearby states (in this case, other states in the New England region).

*Combining agricultural land use intensity with other human impacts*

To create a comprehensive layer of human land use intensity across CONUS, we combined our novel agriculture $L$ layer with an existing $L$ model (CSP 2019) that integrates multiple land use variables into three human impact categories - urban (including data on residential development and nighttime lights), transportation (including roads, railways, powerlines, and pipelines), and energy (including oil and gas wells, coal mines, and utility-scale solar and wind installations) - with each category stored as a separate raster layer. For full details on the existing $L$ layers, including dataset selection and assignment of land use intensity values, see CSP (2019). We combined these three layers with the new agriculture layer into a single land use intensity surface describing the impact at a given location ($L_{loc}$; ranging between 0 and 1) using the “fuzzy algebraic sum" (Theobald 2013). This algorithm ensured that the combined value for a given pixel was always at least as high as that of the most intense disturbance type, but that pixel values never exceeded 1. The fuzzy algebraic sum is given by

$$L_{loc} = 1 - \prod_{j=1}^{k} \left(1 - L_j\right),$$

where $L_j$ is the land use intensity for a given land use type $j$ (i.e., urban, transportation, energy, or agriculture), where $j = 1 \ldots k$ disturbance types (Theobald 2013).

Finally, we incorporated the impact of nearby land uses and disturbances on a given location (Ries *et al.* 2004) by allowing the value of each pixel in our $L_{loc}$ surface to extend beyond the focal pixel itself. We created a new surface of 'neighborhood' land use intensity, $L_n$, by allowing each pixel's $L_{loc}$ value to decay with distance, halving every 500 m out to a maximum distance of 10 km (Theobald *et al.* 2012; CSP 2019). We then combined $L_{loc}$ and $L_n$ to produce our final land use intensity layer ($L_{all}$), again using the fuzzy algebraic sum, i.e.,

$$L_{all} = 1 - ((1 - L_{loc}) * (1 - L_n)).$$

*Summarizing current flow values across regions and land cover/land use types*

We summarized cumulative current flow values from the Omniscape model within regions defined by the USDA Agricultural Research Service (ARS), providing an overview of large-scale differences in agricultural land contributions to connectivity across the country. We calculated the mean and standard deviation of current flow across all agricultural land pixels within each region and also estimated the overall importance of agricultural lands to connectivity in the region. For the latter, we first calculated the upper quartile (i.e., top 25%) value of current flow for all pixels in a given region, regardless of land cover/use type, and then calculated the proportion of those top current flow pixels that occurred on agricultural lands.

We compared current flow on agricultural lands with current flow on other land cover/land use types by first sampling current flow values at > 385,000 random points distributed across CONUS. We classified each random point as falling into one or more of the following categories: *cropland*, *pasture*, *rangeland*, *woodland,* or *all agriculture* (i.e., any one of the previous four categories), based on the FUT 2016 land cover layer (CSP 2020); *low density development,* based on the 2016 National Land Cover Data Base (NLCD; Dewitz 2019) (NLCD classes: 'developed, open space' and 'developed, low intensity' categories); *high density development* (NLCD classes: 'developed, medium intensity' and 'developed, high intensity' categories); *natural land cover* (all NLCD non-agricultural vegetation categories, i.e., cover classes 41-74 and 90-95); and *protected areas* (all public lands in the USGS Protected Areas of the US Database v2.1(USGS 2020) categorized as GAP status 1 or 2, i.e., permanently protected and managed for natural land cover). Agricultural, developed, and natural land categories were mutually exclusive, and we gave preference to FUT agricultural land cover classes where these overlapped with low density development or natural lands. Other categories were non-exclusive; for instance, some natural land pixels were also in protected areas and vice versa. Random point sampling and extraction were conducted in GEE.

*Spatial error regression of the effect of surrounding land cover on agricultural land current flow*

To further explore the drivers of high or low connectivity values on agricultural lands, we also estimate the total amount of natural land cover and development (low and high density development categories combined) within a 1-km radius of each location on agricultural lands, hypothesizing that agricultural lands surrounded by greater amounts of natural land cover and lower levels of development would tend to have higher current flow. This was done for 40,000 randomly selected points on agricultural lands across CONUS (10,000 each for cropland, pasture, rangeland, and woodland). We tested the effect of surrounding land cover/land use on ag land current flow using spatial error regression (Dale and Fortin 2014), modeling current flow as a function of the amount of natural land within 1 km, the amount of developed land within 1 km, and second degree polynomial terms for amounts of natural and developed land to accommodate non-linearity in the response of current flow values to local land cover/use. We also fit a term for agricultural land cover type (categorical: crop, pasture, range, woodland) as well as terms for the interaction between agriculture type and the linear and polynomial effects of the amount of natural land and developed land within 1 km. All predictor variables were mean centered and continuous variables were scaled by one standard deviation prior to model fitting. Current flow values were square root transformed to normalize the spread of data. Prior to model fitting, we confirmed that there was limited correlation between continuous covariates (Pearson's correlation coefficient for amount of natural and developed land: $r = 0.11$). We defined spatial neighbors between the randomly sampled ag land points via Delauney triangulation and calculated a spatial weights matrix using row standardization (Bivand *et al.* 2013). We fit spatial error models using the simultaneous autoregression (SAR) approach (Dale and Fortin 2014) and tested for remaining spatial autocorrelation in the SAR model using Moran's I. Spatial error models were fitted using the *spdep* and *spatialreg* packages in R (R Core Team 2021). In addition to a full model including all terms described above, we fit seven reduced models using subsets of the above terms as well as a null (intercept only) model (nine models total; see Appendix D, Table D1). We compared all models using a model selection approach and Akiake's Information Criterion (AIC; Burnham and Anderson 2002).

*Agricultural land productivity, versatility, and resilience (PVR)*

Agricultural land productivity, versatility, and resilience (PVR) is a CONUS-wide, 10-m resolution data layer describing the long-term sustainability of maintaining a given area in cultivation or other forms of production and is based on soil and land cover characteristics and the type of agriculture practiced at a given location circa 2016. Though technically a snapshot in time, PVR explicitly considers the potential for future disruptions of existing food production systems, thus identifying the "best" agricultural lands for long-term food security. Full details on the calculation of PVR are provided in the peer-reviewed technical documentation accompanying the Farms Under Threat analysis (CSP 2020). Briefly, PVR was calculated as the weighted sum of several (standardized) indicator layers representing soil productivity, land cover and use, food production for direct human consumption, and growing season length. The indicators included and their assigned weights were determined through formal expert elicitation in which 33 agriculture experts from across the U.S. participated in a structured process based on decision analysis theory (Saaty 2008). The values of the resulting PVR layer ranged between 0 and 1 (CSP 2020).

### Appendix B. Validating the relationship between vegetation index variability and management intensity

Previous studies have found that time series of Normalized Difference Vegetation Index (NDVI) data are strongly related to management intensity in agricultural systems subject to frequent harvest/mowing and fertilizer inputs (Franke *et al.* 2012; Gómez Giménez *et al.* 2017). To test whether this relationship holds for agricultural lands across the conterminous U.S., we compared our NDVI coefficient of variation metric (cvNDVI, see main text) (1) between agricultural land cover types; (2) within a given land cover type across a gradient of nitrogen fertilizer input; and (3) between irrigated, unirrigated, and fallow cropland. We began by recalculating cvNDVI for a single year (rather than the five-year timespan used in the full analysis) to better match cvNDVI estimates to the particular crop type and fertilization and irrigation regimes used in a given year. For this validation analysis, we chose the year 2015 to match available fertilization and irrigation datasets (described below). We calculated 2015 cvNDVI for all agricultural cover types (not just cropland and pasture as in the full analysis) to allow comparison between cover types. For each of the four cover types, we randomly selected 20,000 locations across CONUS and extracted 2015 cvNDVI values at each point. For cropland and pasture points, we also extracted estimates of nitrogen fertilizer use in the year 2015 from a 5-km resolution layer developed by Cao et al. (2018). For cropland points, we extracted the crop type planted in 2015 based on the USDA's Cropland Data Layer (CDL) for that year, as well as a binary indicator of whether each cropland point was irrigated or not using a 30-m resolution dataset on irrigation extent across CONUS in the year 2015 (Xie and Lark 2021).

We visually compared the range of cvNDVI values between the four agricultural cover types using density plots, hypothesizing that cropland would exhibit a broader range of values than the other cover types given the potential for greater management intensity on croplands through irrigation, fertilization, multiple crop cycles, etc. Focusing on just cropland and pasture points, we used a spatial regression model (Dale and Fortin 2014) in an ANCOVA framework to test for effects of cover type (crop or pasture), amount of nitrogen fertilizer used, and their interaction on cvNDVI while accounting for spatial autocorrelation arising from proximity of randomly selected locations. Data on nitrogen fertilizer use were mean centered prior to model fitting. We defined spatial neighbors between the randomly sampled points using Delauney triangulation and calculated a spatial weights matrix using row standardization (Bivand *et al.* 2013). We fit spatial error models using the simultaneous autoregression (SAR) approach (Dale and Fortin 2014) and tested for remaining spatial autocorrelation in the SAR model using Moran's I. Spatial error models were fitted using the *spdep* and *spatialreg* packages in R (R Core Team 2021).

Finally, examined the relationship between crop management intensity and cvNDVI for cropland points. For this analysis, we subsetted our dataset to just those cropland points that were planted as corn, soybeans, or wheat in 2015 (the three most common crop types in our dataset) or were left fallow, as determined by the 2015 CDL. We coded each point as falling into one of three management intensity categories - fallow, unirrigated crop, and irrigated crop - and again fit a spatial error model (using the procedure just described) testing the effect of cropland management intensity on cvNDVI. We compared means and 95% confidence interval (CI) values of cvNDVI from the spatial error model between each of the three management intensity categories.

We found that our estimate of vegetation cover variability (cvNDVI) performed well as a proxy for agricultural land management intensity. Across all randomly selected points, cropland, the cover type typically associated with the highest level of anthropogenic activity, had cvNDVI values that tended to be higher and more variable than those of all other agricultural cover types (Fig. B1).

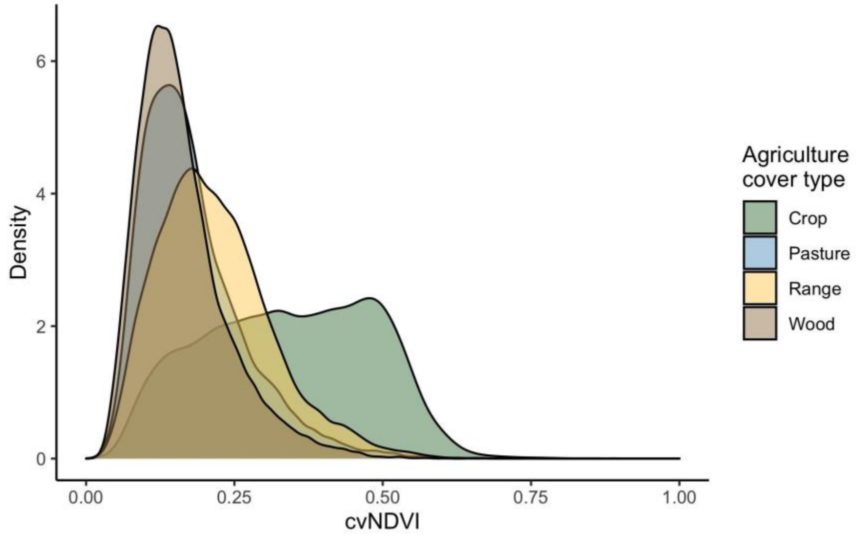

**Figure B1**. Density plot of the distribution of vegetation cover variability (estimated as the coefficient of variation of Normalized Difference Vegetation Index values, or cvNDVI) across all random points for each agricultural cover type.

The spatial error SAR model on nitrogen fertilizer and land cover type sufficiently accounted for spatial autocorrelation in the residuals (Moran's I = -0.05, $p$ = 0.99). The model revealed that cropland has on average a significantly higher cvNDVI than pasture and that, for both cover types, cvNDVI is positively related to nitrogen fertilizer usage but increases more quickly with fertilizer amount on cropland than on pasture (Fig. B2, Table B1).

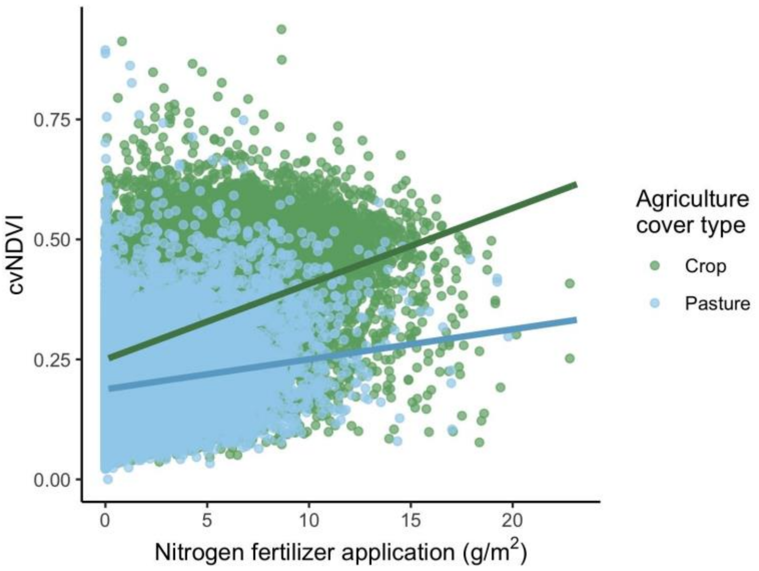

**Figure B2**. Relationship between nitrogen fertilizer usage and variation in vegetation cover (cvNDVI) on cropland (green) and pastureland (blue). Trend lines are fitted values from a spatial error regression model. For both agricultural cover types, increasing fertilizer usage is positively associated with cvNDVI, but with higher slope for cropland (Table 1).

**Table B1.** Spatial error regression analysis results for the effects of agricultural land cover type and nitrogen fertilizer use on variation in vegetation cover (cvNDVI).

| | **Estimate** | **SE** | **P-value** |
|---|---|---|---|
| Intercept (Cover = Crop, Fertilizer = 0) | 0.2996 | 0.0013 | < 0.001 |
| Cover type (Pasture) | 0.0158 | 0.0003 | < 0.001 |
| Fertilizer | -0.0921 | 0.0011 | < 0.001 |
| Cover type x Fertilizer | -0.0095 | 0.0004 | < 0.001 |

The spatial error SAR model on cropland management intensity sufficiently accounted for spatial autocorrelation in the residuals (Moran's I = -0.07, $p$ = 0.99) and revealed that more intensively managed cropland locations exhibited higher cvNDVI. Fallow fields had the lowest average cvNDVI, followed by unirrigated crops, with irrigated crops having the highest cvNDVI. Ninety-five percent confidence intervals, estimated by the spatial error model, did not overlap between management intensity categories (Fig. B3).

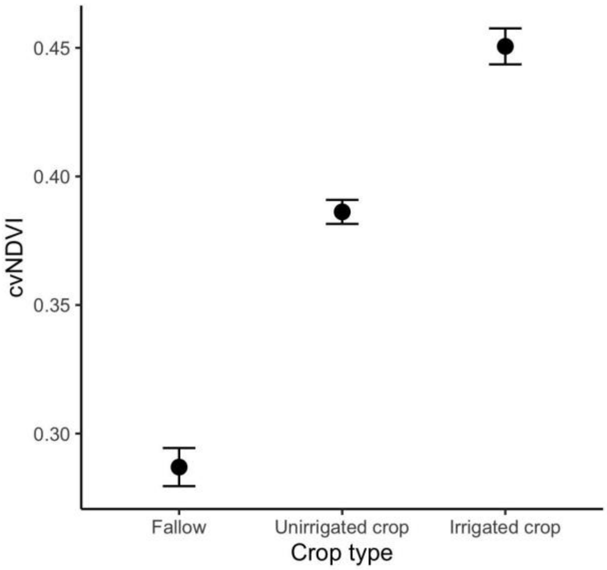

**Figure B3.** Mean (± 95% CI) values of vegetation cover variability (cvNDVI) for fallow agricultural fields, unirrigated crops, and irrigated crops. Means and CIs were calculated from a spatial error regression model.

**Appendix C. Sensitivity of connectivity model results to resistance surface scaling and moving window size.**

Connectivity model results are driven by the relative resistance values applied to different land cover types (Zeller *et al.* 2012). In the case of structural connectivity models based on estimates of the degree or intensity of human land use at each location across the study area, the scaling function used to convert land use intensity to resistance is therefore a key consideration. For the analyses presented in the main text, we followed Dickson et al. (2017) in calculating resistance ($R$) from human land use intensity ($L$) as

$$R = (L + 1)^{10} + s/4,$$

where $s$ is the percent slope of a given pixel (see main text for additional discussion and justification of this scaling function). The above rescaling formula led to a relatively high contrast between the resistance values assigned to locations with low, medium, and high $L$ values (Dickson *et al.* 2017). It was not our goal here to conduct a comprehensive sensitivity analysis of the effects of different scaling functions on resistance and connectivity; this analysis has already been conducted by Dickson et al. (2017) using a human land use intensity layer derived via very similar methods to those employed here. For comparison, however, we derived a second resistance surface using a low-contrast rescaling formula suggested by Marrec et al. (2020):

$$R = 1 + (1000 * L^2) + (s/4).$$

We then used this new, low-contrast resistance surface to fit a connectivity model in Omniscape, keeping all other model inputs and parameters (e.g., moving window radius, source strength surface) the same as those of the model presented in the main text.

The maximum distance between current flow start and end points is another key assumption of circuit theory-based connectivity models, controlled in Omniscape models by the radius of the moving window used to calculate current flow between source and target locations (McRae *et al.* 2016; Landau *et al.* 2021). In connectivity analyses focused on particular species or guilds, this value is often set to the maximum (or average) movement or dispersal distance of that species or guild (e.g., Littlefield *et al.* 2017; Jennings *et al.* 2020). Here we were interested in modeling potential connectivity for a wide range of species as an estimate of ecological flow. We therefore chose to use a moving window radius comparable the upper dispersal distances of many large-bodied terrestrial vertebrates (Sutherland *et al.* 2000) under the assumption that landscapes capable of supporting the movements of species with large space requirements will also support the movement of less vagile species. However, for comparison, we also fit connectivity models using two smaller moving window radii, (i) 20 km, comparable to the maximum dispersal distance of many medium sized mammals (Sutherland *et al.* 2000), and (ii) 5 km, approximating maximum dispersal distances for small vertebrates such as amphibians; (Marsh and Trenham 2001). For the connectivity models with smaller moving window radii, we kept all model inputs (i.e., resistance and source strength surfaces) the same as for the model presented in the main text, but varied the block size parameter to better match the moving window sizes used. Omniscape block size controls the density of pixels in a landscape raster that can potentially be treated as targets for current flow and can be used to decrease computation time with large landscape rasters (Landau *et al.* 2021). If block size = 1, all pixels are treated as potential targets (as long as source strength at the pixel is nonzero), with block size = 3, every third pixel is a potential target, and so on. We set block size to 21 for the 20-km model and 3 for the 5-km model. All models described here were based on 250-m resolution rasters.

To test the robustness of our conclusions regarding the relationship between current flow and land cover/use to model assumptions, we used the procedure described in the main text to extract current flow values for our primary connectivity model (i.e., the one presented in the main text) and for each of the three comparison models (i.e., the low-contrast, 20-km, and 5-km models) at > 385,000 random points distributed across the conterminous US (CONUS). Each point was assigned to one or more of the following land cover/use categories: cropland, pasture, rangeland, woodland, all agriculture, low density development, high density development, natural land cover, and protected areas (see main text for details). For each land cover/use category, we calculated Spearman's rank correlations (ρ) between current flow values from the primary model and those from each of the three comparison models. We also calculated the average current flow value for each land cover/use category under each model and compared the rank order of current flow values across categories between the primary model and each of the comparison models using Spearman's rank correlations.

Correlation coefficients for each comparison are shown in Table C1 and suggest that model assumptions regarding the scaling function used to create the resistance surface and the size of the Omniscape moving window have relatively limited effects on the relationship between land cover/use and current flow. For all comparison models, land cover rankings were highly correlated with those of the primary model (ρ ≥ 0.99). For individual land cover classes, current flow values derived from the primary model were typically strongly correlated with those derived from each of the three comparison models (Table C1), with the weakest correlations being for the high-density development class with the 20-km (ρ = 0.68) and 5-km (ρ = 0.59) models.

**Table C1.** Spearman's rank correlations (ρ) between current flow values derived from the primary connectivity model (i.e., the model presented in the main text) and each of the three comparison models (described above) for each land cover/use category. The bottom row presents correlations for the rank order of average current flow values for each land cover/use category.

| **Land cover/use category** | **Low-contrast scaling, ρ** | **20 km moving window, ρ** | **5 km moving window, ρ** |
|---|---|---|---|
| Cropland | 0.89 | 0.89 | 0.84 |
| Pasture | 0.93 | 0.91 | 0.86 |
| Rangeland | 0.92 | 0.88 | 0.80 |
| Woodland | 0.95 | 0.92 | 0.87 |
| All agriculture | 0.95 | 0.95 | 0.92 |
| Development, low density | 0.94 | 0.87 | 0.83 |
| Development, high density | 0.91 | 0.68 | 0.59 |
| Natural | 0.92 | 0.89 | 0.82 |
| Protected areas | 0.91 | 0.83 | 0.71 |
| *All categories, ranked* | 0.99 | 0.99 | 1.00 |

Finally, we mapped current flow across the CONUS for each of the three comparison models (Fig. C1), highlighting differences between models in the relative intensity and concentration of current flow. Current flow tended to be more diffuse in the 20-km and 5-km models relative to the low-contrast model (which used a 150-km moving window radius), reflecting the shorter movement distances allowed in the 20-km and 5-km models.

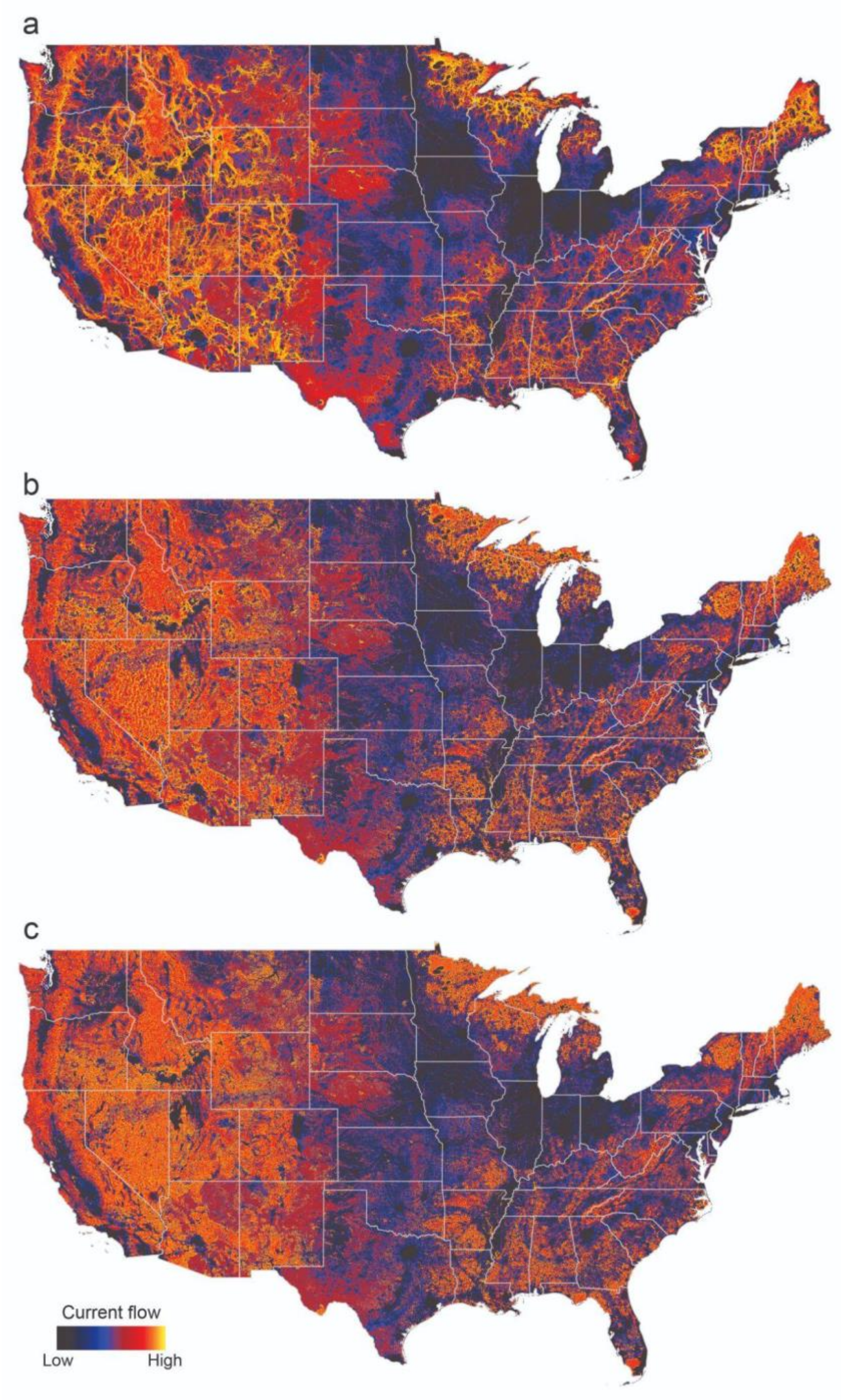

**Figure C1.** Current flow maps derived from each of the three comparison connectivity models described in Appendix C: (**a**) the low-contrast model, (**b**) the 20-km model, and (**c**) the 5-km model. Current flow is proportional to the potential net movement of organisms through a given location on the landscape.

### Appendix D. Supplementary tables and figures

**Table D1.** Model selection table for the spatial error regression model describing the effects of surrounding land cover/land use on agricultural land connectivity. The response variable in all models was the square root of current flow sampled at 40,000 random locations on agricultural lands across the conterminous United States. *nat* = amount of natural land within 1 km of the sampled location. *dev* = amount of developed land within 1 km of the sampled location. *ag type* = agricultural land cover class (cropland, pasture, rangeland, or woodland, as defined in the main text). Developed and natural land cover/use categories are based on the 2016 Nation Land Cover Database and defined in the main text.

| **Model formula** | **Degrees of freedom** | **AIC** | **ΔAIC** |
|---|---|---|---|
| nat + $nat^2$ + dev + $dev^2$ + ag type + nat * ag type + $nat^2$ * ag type + dev * ag type + $dev^2$ * ag type | 22.0 | 194303.3 | 0.0 |
| nat + dev + $dev^2$ + ag type + nat * ag type + dev * ag type + $dev^2$ * ag type | 18.0 | 194417.5 | 114.3 |
| nat + $nat^2$ + dev + $dev^2$ + ag type + dev * ag type + $dev^2$ * ag type | 16.0 | 194459.0 | 155.7 |
| nat + $nat^2$ + dev + $dev^2$ + ag type + nat * ag type + $nat^2$ * ag type | 16.0 | 194716.0 | 412.8 |
| nat + $nat^2$ + dev + ag type + nat * ag type + $nat^2$ * ag type + dev * ag type | 18.0 | 194738.0 | 434.7 |
| nat + dev + ag type + nat * ag type + dev * ag type | 14.0 | 194880.9 | 577.7 |
| nat + $nat^2$ + dev + $dev^2$ + ag type | 10.0 | 194935.2 | 632.0 |
| nat + dev + ag type | 8.0 | 195342.7 | 1039.5 |
| null model (~ 1) | 3.0 | 207905.0 | 13601.7 |

**Table D2.** Summary of landscape resistance and current flow values on agricultural lands (cropland, pasture, rangeland, and woodland, as well as all agricultural categories combined) compared to values in developed areas and natural landscapes. Developed and natural land cover/use categories are based on the 2016 Nation Land Cover Database and defined in the main text. Protected areas are USGS GAP 1 and 2 protected areas. Resistance and current flow values are presented as means (standard deviations).

| Land cover and use category | Resistance | Current flow |
|---|---|---|
| Cropland | 372.1 (202.1) | 91.1 (56.0) |
| Pasture | 217.9 (168.1) | 133.4 (84.4) |
| Rangeland | 65.1 (101.2) | 226.6 (92.9) |
| Woodland | 149.1 (152.3) | 169.6 (105.9) |
| All agriculture | 208.7 (211.2) | 158.8 (101.5) |
| Development, low density | 324.6 (247.6) | 131.2 (147) |
| Development, high density | 490 (202.5) | 59.5 (53.3) |
| Natural | 39.9 (112.3) | 378.5 (196.6) |
| Protected areas | 44.5 (161.7) | 401.2 (194.8) |

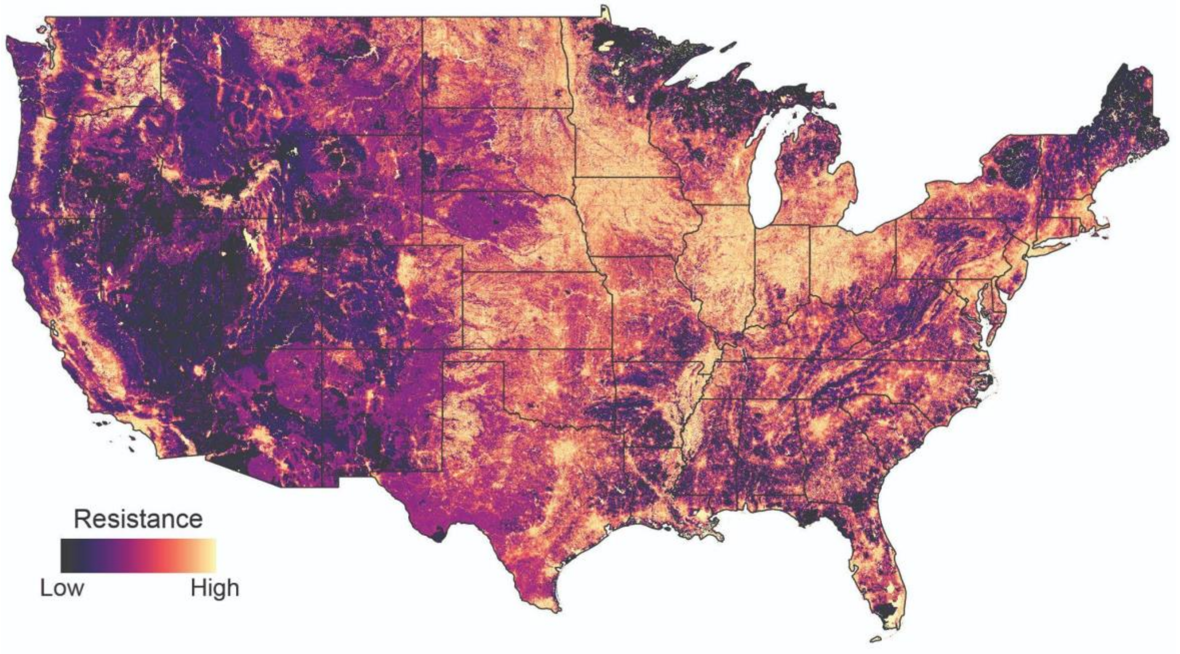

**Figure D1.** Map of landscape resistance across the conterminous United States

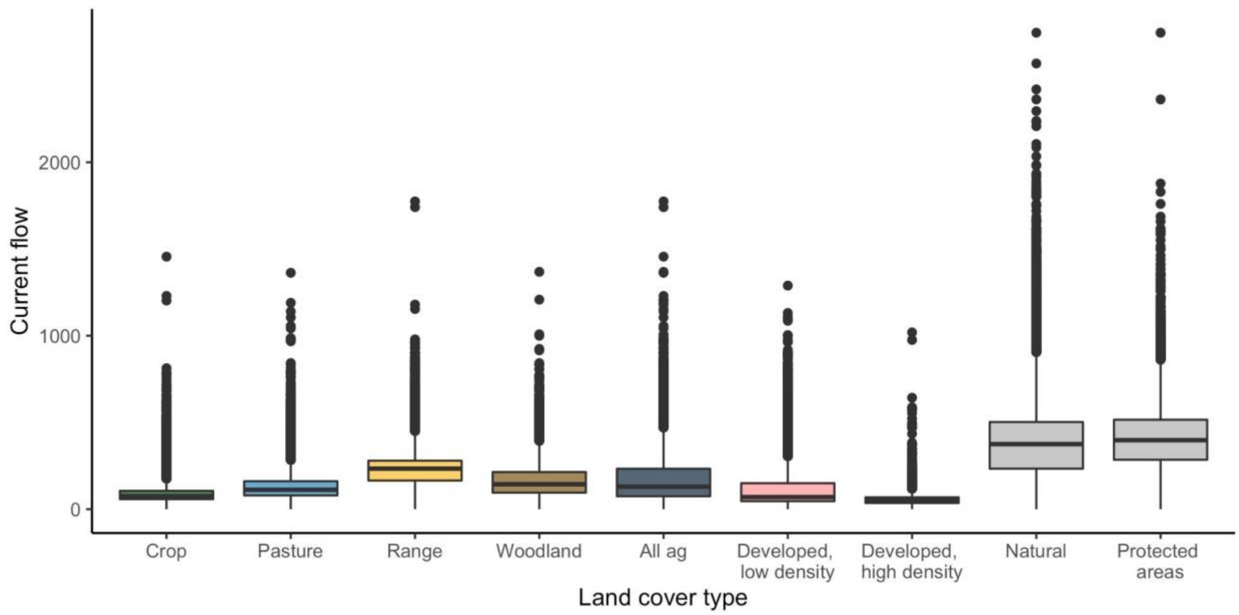

**Figure D2.** The range of current flow on agricultural lands (cropland, pasture, rangeland, and woodland, as well as all agricultural categories combined [‘all ag’]) is compared to that of developed areas and landscapes characterized by more natural land cover types (i.e., all natural lands and those within USGS GAP 1 or GAP 2 protected areas). Data are shown as standard boxplots with whiskers representing 1.5 times the interquartile range. Outliers are shown as points.

### Supplementary References

Bivand RS, Pebesma E, and Gómez-Rubio V. 2013. Applied Spatial Data Analysis with R. New York: Springer.

Burnham KP and Anderson DR. 2002. Model Selection and Multimodel Inference: A Practical Information-Theoretic Approach. New York: Springer.

Cao P, Lu C, and Yu Z. 2018. Historical nitrogen fertilizer use in agricultural ecosystems of the contiguous United States during 1850–2015: application rate, timing, and fertilizer types. *Earth System Science Data* **10**: 969–84.

CSP. 2019. Methods and approach used to estimate the loss and fragmentation of natural lands in the conterminous U.S. from 2001 to 2017. Truckee, CA.

CSP. 2020. Description of the approach, data, and analytical methods used for the Farms Under Threat: State of the States project, version 2.0. Final Technical Report. Truckee, CA.

Dale MRT and Fortin M-J. 2014. Spatial Analysis: A Guide For Ecologists. Cambridge University Press.

Dewitz J. 2019. National Land Cover Database (NLCD) 2016 Products: U.S. Geological Survey data release.

Dickson BG, Albano CM, McRae BH, *et al.* 2017. Informing strategic efforts to expand and connect protected areas using a model of ecological flow, with application to the western United States. *Conservation Letters* **10**: 564–71.

Franke J, Keuck V, and Siegert F. 2012. Assessment of grassland use intensity by remote sensing to support conservation schemes. *Journal for Nature Conservation* **20**: 125–34.

Gómez Giménez M, Jong R de, Della Peruta R, *et al.* 2017. Determination of grassland use intensity based on multi-temporal remote sensing data and ecological indicators. *Remote Sensing of Environment* **198**: 126–39.

Jennings MK, Zeller KA, and Lewison RL. 2020. Supporting Adaptive Connectivity in Dynamic Landscapes. *Land* **9**: 295.

Landau VA, Shah VB, Anantharaman R, and Hall KR. 2021. Omniscape.jl: Software to compute omnidirectional landscape connectivity. *Journal of Open Source Software* **6**: 2829.

Littlefield CE, McRae BH, Michalak JL, *et al.* 2017. Connecting today's climates to future climate analogs to facilitate movement of species under climate change. *Conservation Biology* **31**: 1397–408.

Marrec R, Abdel Moniem HE, Iravani M, *et al.* 2020. Conceptual framework and uncertainty analysis for large-scale, species-agnostic modelling of landscape connectivity across Alberta, Canada. *Scientific Reports* **10**: 6798.

Marsh DM and Trenham PC. 2001. Metapopulation Dynamics and Amphibian Conservation. *Conservation Biology* **15**: 40–9.

McRae B, Popper K, Jones A, *et al.* 2016. Conserving nature's stage: Mapping omnidirectional connectivity for resilient terrestrial landscapes in the Pacific Northwest. The Nature Conservancy, Portland, OR.

R Core Team. 2021. R: A language and environment for statistical computing. Vienna, Austria: R Foundation for Statistical Computing.

Ries L, Fletcher RJ, Battin J, and Sisk TD. 2004. Ecological responses to habitat edges: Mechanisms, models, and variability explained. *Annual Review of Ecology, Evolution, and Systematics* **35**: 491–522.

Saaty TL. 2008. Decision making with the analytic hierarchy process. *International Journal of Services Sciences* **1**: 83–98.

Sacks WJ, Deryng D, Foley JA, and Ramankutty N. 2010. Crop planting dates: an analysis of global patterns. *Global Ecology and Biogeography* **19**: 607–20.

Sutherland GD, Harestad AS, Price K, and Lertzman KP. 2000. Scaling of Natal Dispersal Distances in Terrestrial Birds and Mammals. *Conservation Ecology* **4**.

Theobald DM, Reed SE, Fields K, and Soulé M. 2012. Connecting natural landscapes using a landscape permeability model to prioritize conservation activities in the United States. *Conservation Letters* **5**: 123–33.

USGS. 2020. U.S Geological Survey (USGS) Gap Analysis Project (GAP). Protected Areas Database of the United States (PAD-US) 2.1: U.S. Geological Survey data release.

Xie Y and Lark TJ. 2021. Mapping annual irrigation from Landsat imagery and environmental variables across the conterminous United States. *Remote Sensing of Environment* **260**: 112445.

Zeller KA, McGarigal K, and Whiteley AR. 2012. Estimating landscape resistance to movement: a review. *Landscape Ecol* **27**: 777–97.
